# Gene III® L-Ergothioneine, alone or combined with vitamin K2, vitamin D3 and magnesium L-threonate, attenuates bone turnover, inflammatory and oxidative disturbances in ovariectomized mice

**DOI:** 10.64898/2026.06.23.734114

**Authors:** Wei Liu, Yongxian Tang, Wei Ding, Juan Cao, Cong Guo, Guohua Xiao

## Abstract

**Purpose:** Estrogen deficiency drives bone loss through a complex interplay of endocrine, oxidative, inflammatory, and bone-remodeling disturbances. Ergothioneine (EGT) is a diet-derived thiol/thione antioxidant, yet its effects on the estrogen-deficient skeleton remain unknown. This study evaluated whether EGT–alone or combined with vitamin K2, vitamin D3, and magnesium L-threonate–attenuates the skeletal and systemic consequences of ovariectomy (OVX) in mice.

**Methods:** Forty-eight female C57BL/6J mice underwent either sham surgery or OVX. For 12 weeks, the mice received a daily oral gavage of one of the following: vehicle, alendronate (1.53 mg/kg), EGT (30 mg/kg/day), EGT with vitamin K2 (40 µg/kg/day) and vitamin D3 (500 IU/kg/day), or EGT with vitamin K2, magnesium L-threonate (350 mg/kg/day), and vitamin D3 (n = 5–6 per group). Outcomes evaluated included the uterine index, tibial micro-computed tomography (micro-CT), distal-femoral histology, serum bone turnover markers (CTX-I, PINP, osteocalcin), sex hormones, TNF-α, IL-6, SOD, and MDA.

**Results:** OVX significantly lowered the uterine index and induced tibial trabecular deterioration. This was accompanied by increased CTX-I, TNF-α, IL-6, and MDA, alongside decreased PINP, osteocalcin, and SOD (all P < 0.01 vs. sham). Alendronate successfully restored tibial micro-CT bone-volume fraction (BV/TV) and trabecular number (P < 0.01 vs. OVX). While the EGT-based regimens did not significantly restore tibial micro-CT BV/TV, trabecular thickness, or trabecular number (all P > 0.05 vs. OVX), they significantly increased the trabecular area in distal-femoral histology (14.2–15.0% across regimens vs. 7.6% in OVX; P < 0.05). Furthermore, EGT treatments lowered CTX-I, TNF-α, IL-6, and MDA, increased SOD, and partially restored PINP and osteocalcin (P < 0.05–0.01 vs. OVX). Apparent increases in serum estradiol were assay-dependent and are considered exploratory.

**Conclusion:** EGT-based nutritional regimens improved the systemic oxidative, inflammatory, and bone-turnover environments associated with estrogen-deficient bone loss and successfully preserved the distal-femoral trabecular area. However, tibial three-dimensional microarchitecture was not restored on micro-CT. Given that histological and micro-CT endpoints were evaluated at different skeletal sites, structural interpretations should be made with caution. These findings support the further evaluation of EGT as a dietary adjunct, warranting subsequent mechanistic and dose-optimization studies.

## Introduction

Postmenopausal osteoporosis arises chiefly from estrogen deficiency and presents with reduced bone mass, disrupted trabecular architecture, and increased fracture risk (Compston et al. 2019; Eastell et al. 2016). Its pathophysiology extends beyond the loss of sex-steroid signaling: estrogen withdrawal raises osteoclastogenic inflammatory cytokines, promotes oxidative stress, impairs osteoblast function, and tips bone remodeling toward net resorption (Khosla et al. 2012; Lean et al. 2003; Manolagas 2010; Wauquier et al. 2009; Weitzmann and Pacifici 2006). Ovariectomized rodents recapitulate many of these features and remain a standard model for preclinical testing of anti-osteoporotic interventions (Komori 2015; Lelovas et al. 2008).

Because inflammation and oxidative stress are central to this process, they are attractive intervention targets. TNF-α and IL-6 drive osteoclast differentiation and activity, whereas oxidative stress suppresses osteoblastogenesis and hastens osteoblast and osteocyte dysfunction (Khosla et al. 2012; Lean et al. 2003; Manolagas 2010; Wauquier et al. 2009; Weitzmann and Pacifici 2006). Circulating markers such as CTX-I, PINP, osteocalcin, SOD, and MDA thus complement structural imaging by capturing the biological state of bone remodeling (Eastell and Szulc 2017; Vasikaran et al. 2011). Agents acting on these convergent pathways may serve as supportive or adjunctive strategies, especially where they improve the remodeling environment before microarchitectural recovery becomes detectable.

Ergothioneine is a sulfur-containing micronutrient taken up via the transporter OCTN1/SLC22A4 and valued for its antioxidant and cytoprotective activity (Cheah and Halliwell 2012; Gründemann et al. 2005; Halliwell et al. 2018). Its capacity to buffer oxidative stress and temper inflammatory signaling makes it a reasonable candidate for estrogen-deficiency-induced bone loss. Vitamin K2 and vitamin D3 likewise participate in skeletal mineral metabolism, and adequate magnesium status is linked to bone health (Bouillon et al. 2019; Castiglioni et al. 2013; Knapen et al. 2013; Rondanelli et al. 2021). Whether EGT, alone or with these nutrients, can preserve trabecular structure, normalize bone turnover, and modify inflammatory or oxidative markers in vivo has not been established.

We therefore examined EGT alone, EGT with vitamin K2 and vitamin D3, and EGT with vitamin K2, magnesium L-threonate, and vitamin D3 in C57BL/6J OVX mice. Uterine index, micro-CT, histology, serum bone turnover markers, sex hormones, inflammatory cytokines, and oxidative stress assays were combined to test whether these regimens attenuate OVX-induced skeletal and systemic pathology.

## Materials and Methods

### Animals

Forty-eight female C57BL/6J mice, 8 weeks old and weighing 19 to 21 g, were obtained from Zhejiang Vital River Laboratory Animal Technology Co., Ltd. The animals were maintained under SPF conditions. The animal certificate number was 20251014Abzz06000000124, and the experimental animal production license was SCXK (Su) 2025-0029. Mice were housed at 20 to 26 °C with daily temperature fluctuation not exceeding 4°C, relative humidity of 40% to 70%, at least 15 air changes per hour, and a 12 h light/dark cycle from 07:00 to 19:00. Standard SPF rodent maintenance diet and purified drinking water were provided ad libitum unless otherwise specified. Animal experiments were approved under ethics approval number I20250006 and are reported in accordance with ARRIVE 2.0 principles (Percie du Sert et al. 2020).

### Experimental Groups and Treatments

After 5 days of acclimation, mice were weighed and randomized into six groups. Forty-eight mice were obtained to provide a margin against perioperative or unexpected loss; the final analyzed datasets contained five or six animals per group depending on endpoint availability. The groups were sham control, OVX model, alendronate positive control, Gene III® L-EGT alone (EGT), Gene III® L-EGT plus vitamin K2 and vitamin D3 (EGT+K2/D3), and Gene III® L-EGT plus vitamin K2, magnesium L-threonate and vitamin D3 (EGT+K2/MgT/D3). The sham and OVX groups received an equivalent volume of water by gavage. The positive-control group received alendronate at 1.53 mg/kg. EGT was given at 30 mg/kg/day. The EGT+K2/D3 group received EGT 30 mg/kg/day, vitamin K2 40 µg/kg/day and vitamin D3 500 IU/kg/day; the EGT+K2/MgT/D3 group received EGT 30 mg/kg/day, vitamin K2 40 µg/kg/day, magnesium L-threonate 350 mg/kg/day and vitamin D3 500 IU/kg/day. These per-kilogram doses correspond to approximate human-equivalent daily intakes of vitamin K2 160 µg/day and vitamin D3 2000 IU/day. Treatments were administered by oral gavage once daily for 12 weeks.

### OVX Model Establishment

Except for the sham control group, mice underwent bilateral ovariectomy. Briefly, after anesthesia and preparation of the surgical field, the skin, fascia, and muscle layers were opened, adipose tissue was separated, and the ovaries were exposed. The oviducts were ligated, and both ovaries were excised. Sham-control mice underwent sham surgery without ovarian removal. The incision was closed layer by layer and disinfected with iodophor. Mice received low-dose penicillin intramuscularly for 3 consecutive days to prevent infection. One week after surgery, OVX mice were fed a low-calcium, low-protein diet for 1 week, after which gavage intervention began.

### Uterine Weight and Organ Index

At the end of the 12-week treatment period, mice were euthanized, and uteri were dissected and weighed using an electronic balance. The uterine organ index was calculated to verify the estrogen-deficiency state induced by OVX. Uterine weight and organ index were summarized as mean ± SD.

### Micro-Computed Tomography

After the final gavage, mice were fasted for 12 h and euthanized. Tibial specimens were dissected free of surrounding muscle and stored at −80 °C before imaging, then scanned on a SkyScan 1276 micro-CT system (CTAn v1.23.0.1; isotropic pixel size ≈ 17.76 µm; grey-scale threshold 45–255). Using CTAn, we quantified bone-volume fraction (BV/TV, %), trabecular thickness (Tb.Th, mm), trabecular number (Tb.N, mm□^1^), trabecular separation (Tb.Sp, mm), trabecular pattern factor (Tb.Pf, mm□^1^), structure model index (SMI), and connectivity density (Conn.D, mm□^3^); three-dimensional reconstructions were generated for visualization. As no mineral-density calibration phantom was scanned, true volumetric bone mineral density was not derived. Micro-CT acquisition and analysis followed established rodent bone-microstructure recommendations (Bouxsein et al. 2010).

### Histological Assessment

Femoral distal metaphyseal bone was evaluated by hematoxylin and eosin staining. Trabecular morphology was assessed at low and high magnification. Trabecular number was estimated using two parallel 1 mm lines separated by 0.2 mm; intersections between the test lines and trabecular bone were counted, and histological Tb.N was calculated as P/ (L × 2). Trabecular area was measured from a 1.5 mm marrow cavity region beginning at the epiphyseal line using ImageJ and expressed as a percentage of the marrow cavity area. Empty lacuna rate was calculated as the number of empty lacunae divided by total lacunae multiplied by 100%. No empty lacunae were observed in this model.

### Serum ELISA and Biochemical Assays

After 12 h fasting at the end of treatment, mice were weighed and anesthetized for blood collection. Serum was separated and stored at -20 °C. ELISA assays were used to measure estradiol (E2), follicle-stimulating hormone (FSH), luteinizing hormone (LH), TNF-α, IL-6, PINP, CTX-I, and osteocalcin/BGP. The ELISA kits were from Shanghai Kexing Trading Co., Ltd., with catalog numbers F30285-A for PINP, F30282-A for CTX-I, F2551-A for osteocalcin/BGP, F2546-A for E2, F2555-A for FSH, and F2582-A for LH. Samples were diluted 5-fold in the ELISA workflow, incubated with HRP-labeled detection antibody at 37 °C for 60 min, developed with substrate A/B for 15 min, stopped, and read at 450 nm.

Serum SOD activity was measured using a WST-8 microplate assay kit (ADS-W-KY011), and MDA was measured using a microplate thiobarbituric acid-related assay kit (ADS-W-YH002). The SOD and MDA assay kits were supplied by Shanghai Addison Biotechnology Co., Ltd. (Shanghai, China). SOD absorbance was read at 450 nm, and MDA absorbance was read at 532 and 600 nm.

### Statistical Analysis

Data are expressed as mean ± SD and were analyzed in GraphPad Prism 10. Homogeneity of variance was assessed by Levene’s test; when variances were homogeneous we used one-way ANOVA, otherwise the Kruskal–Wallis test. After a significant global test, group comparisons used Dunnett’s test or the Mann–Whitney U test. Significance versus the sham control is denoted by asterisks (*P < 0.05, **P < 0.01) and versus the OVX model group by hash symbols (#P < 0.05, ##P < 0.01).

## Results

### Uterine Index and Model Validation

At the end of the experiment, uterine weight and the uterine organ index were markedly lower in OVX than in sham mice, confirming successful estrogen-deficiency modeling. Uterine weight fell from 0.139 ± 0.029 g (sham) to 0.052 ± 0.027 g (OVX), and the uterine index from 0.67 ± 0.13% to 0.24 ± 0.15% (P < 0.01 vs sham). Alendronate and all three EGT-based regimens showed partial numerical recovery, but none differed significantly from the OVX group (P > 0.05).

### Tibial Trabecular Microarchitecture (Micro-CT)

Micro-CT confirmed OVX-associated deterioration of tibial trabecular bone. Bone-volume fraction (BV/TV) decreased from 11.6 ± 1.0% (sham) to 8.8 ± 0.5% in OVX mice (P < 0.01), Tb.Th from 0.074 ± 0.0039 to 0.063 ± 0.0054 mm (P < 0.05), and Tb.N from 1.455 ± 0.378 to 0.708 ± 0.507 mm□^1^ (P < 0.05). Alendronate significantly increased BV/TV and Tb.N relative to OVX, to 12.2 ± 1.6% and 1.972 ± 0.785 mm□^1^ (both P < 0.01) (Fig. 1).

**Figure 1.**
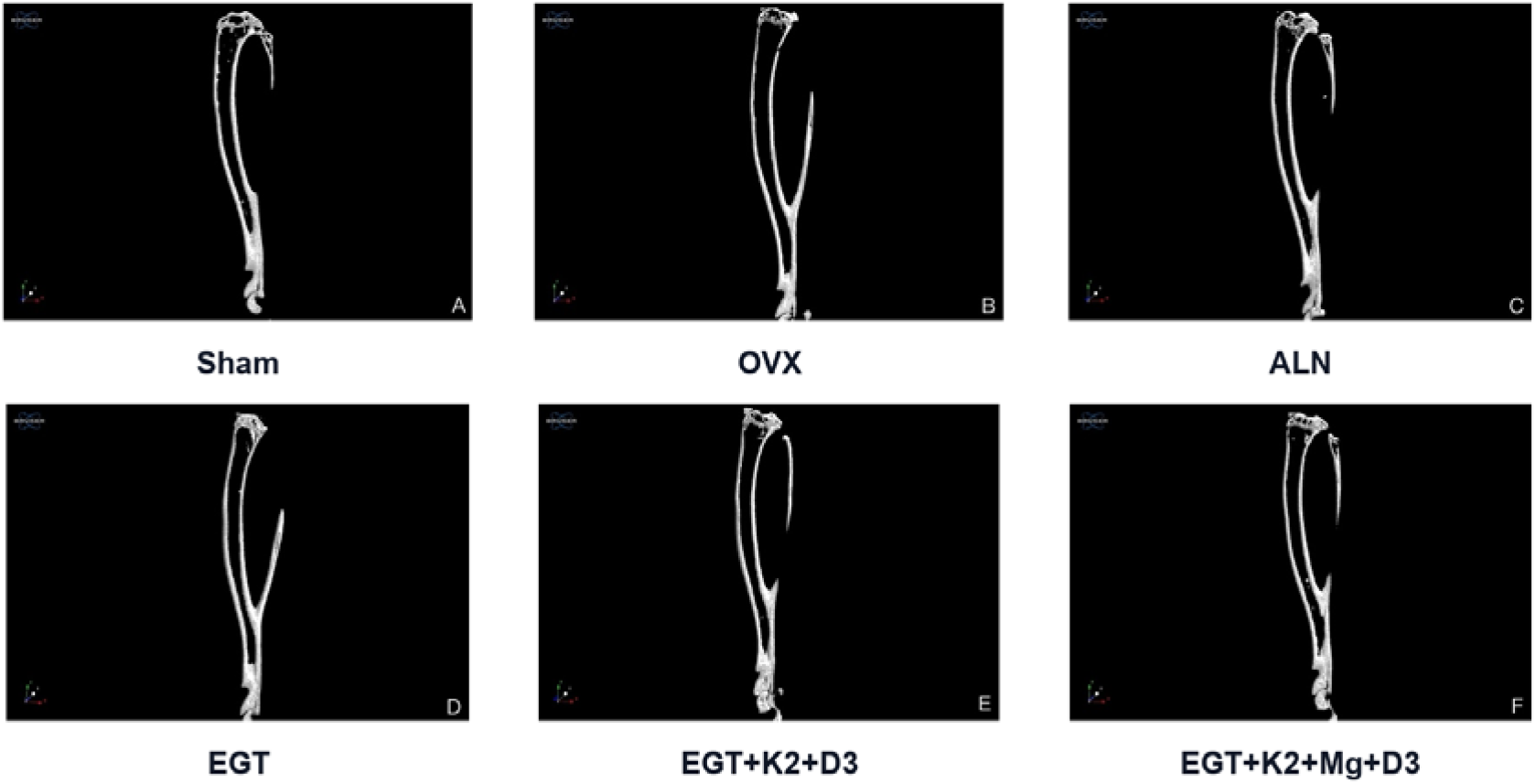
Representative micro-CT images after 12 weeks of treatment. Representative tibial micro-CT reconstructions are shown for sham control (A), OVX model (B), alendronate (C), ergothioneine (D), ergothioneine plus vitamin K2 and vitamin D3 (E), and ergothioneine plus vitamin K2, magnesium L-threonate, and vitamin D3 (F).

The three EGT-based regimens produced only non-significant numerical trends in BV/TV, Tb.Th and Tb.N relative to the OVX group (all P > 0.05). BV/TV was 10.1 ± 0.5% (EGT), 10.0 ± 0.8% (EGT+K2/D3) and 10.0 ± 1.7% (EGT+K2/MgT/D3); Tb.Th was 0.067 ± 0.0058, 0.066 ± 0.0066 and 0.068 ± 0.0088 mm; and Tb.N was 0.976 ± 0.161, 0.978 ± 0.260 and 0.986 ± 0.702 mm□^1^, respectively. Thus the positive control validated the model and assay sensitivity, whereas the EGT-based regimens achieved little three-dimensional structural recovery at the tested doses and duration. Connectivity density in the EGT-based groups was numerically lower than in the OVX group (Supplementary Table S2), indicating that trabecular connectivity was not restored (Fig. 2).

**Figure 2.**
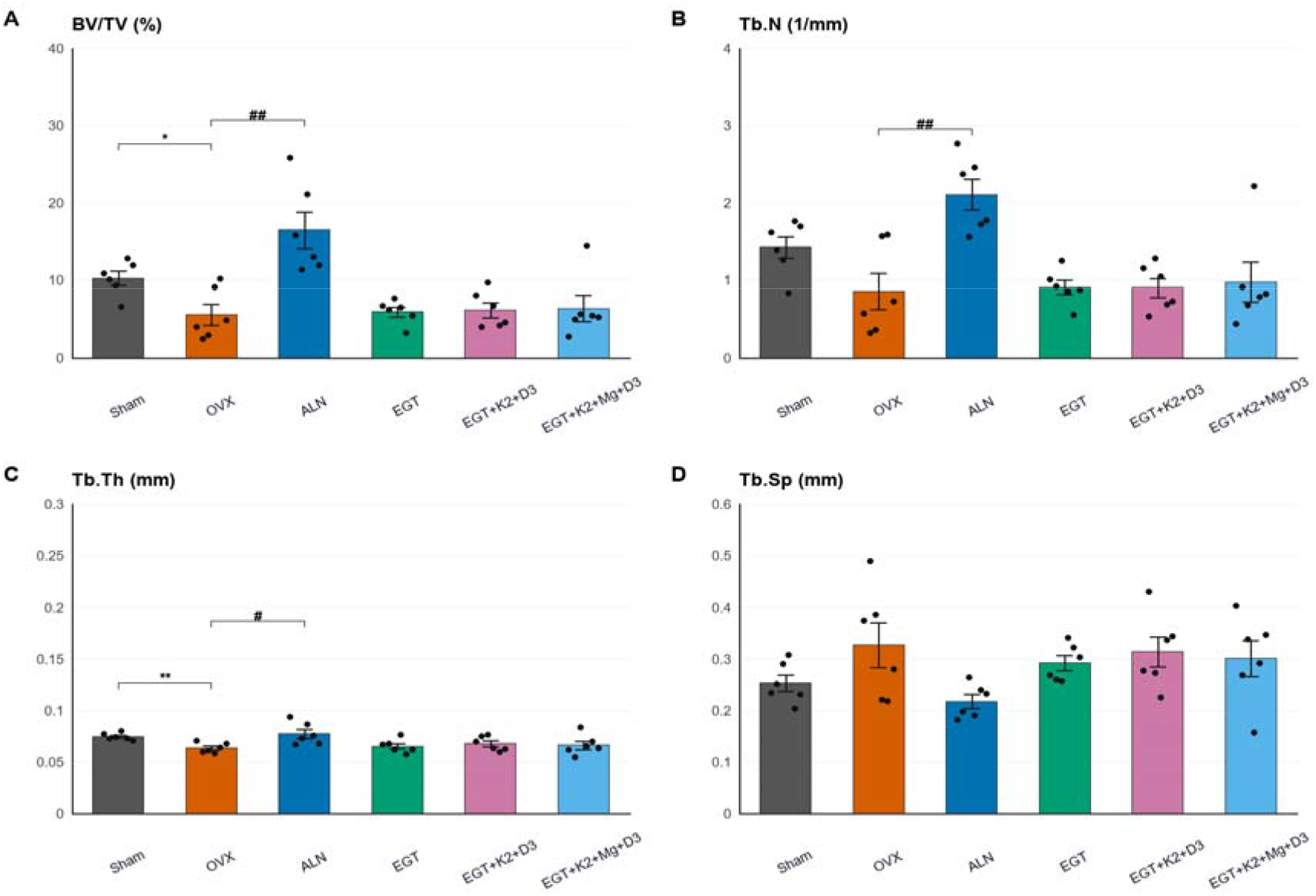
Quantitative micro-CT assessment of trabecular bone. Quantitative micro-CT parameters show OVX-induced bone loss and strong protection by alendronate. Ergothioneine alone and ergothioneine-based combination regimens showed non-significant trends toward improved bone density, trabecular thickness, and trabecular number.

### Distal-Femoral Trabecular Histology

Hematoxylin–eosin staining mirrored the micro-CT findings. Sham controls showed abundant, regular, plate-like to mildly rod-like trabeculae of fairly uniform thickness, whereas OVX mice showed sparse, thin, isolated and partly disrupted trabeculae. Alendronate-treated bone approached the sham appearance, while the EGT and combination groups were intermediate and heterogeneous–some specimens retained more abundant, regular trabeculae and others still showed thinning and loss (Fig. 3).

**Figure 3.**
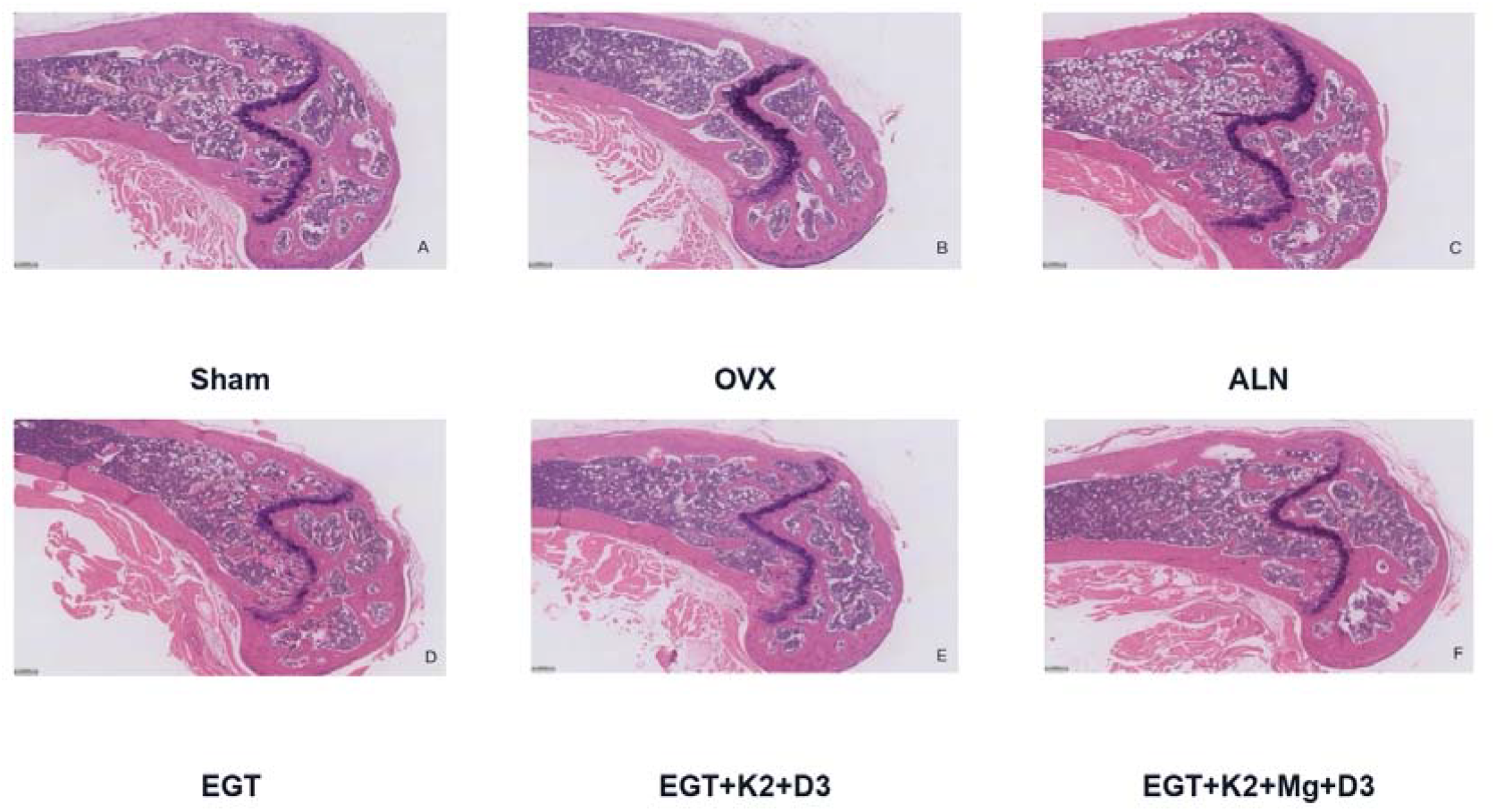
Histological evaluation of distal femoral trabecular bone. HE staining showed abundant regular trabeculae in sham controls, sparse and disrupted trabeculae in OVX mice, near-normal trabecular morphology in alendronate-treated mice, and partial preservation in ergothioneine-treated and combination-treated mice. Quantification showed significant improvement of trabecular area in all ergothioneine-based intervention groups.

Quantitatively, trabecular area on distal-femoral histology was significantly reduced in OVX mice, from 16.41 ± 4.50% (sham) to 7.56 ± 2.62% (P < 0.01). Alendronate restored it to 16.90 ± 1.52% (P < 0.01 vs OVX). EGT, EGT+K2/D3 and EGT+K2/MgT/D3 increased trabecular area to 14.19 ± 4.83%, 14.55 ± 3.55% and 14.95 ± 4.46%, respectively, each significantly higher than OVX (P < 0.05). Histological trabecular number showed a downward trend in OVX and partial numerical improvement after treatment but did not reach significance for any group, and no empty lacunae were observed.

### Serum Bone Turnover Markers

OVX produced a bone-turnover profile of increased resorption and impaired formation. CTX-I rose from 4.189 ± 0.883 ng/mL (sham) to 8.237 ± 0.513 ng/mL (P < 0.01); PINP fell from 19.90 ± 1.51 to 10.72 ± 2.11 ng/mL (P < 0.01); and osteocalcin/BGP fell from 78.56 ± 4.24 to 42.14 ± 8.87 ng/mL (P < 0.01) (Fig. 4).

**Figure 4.**
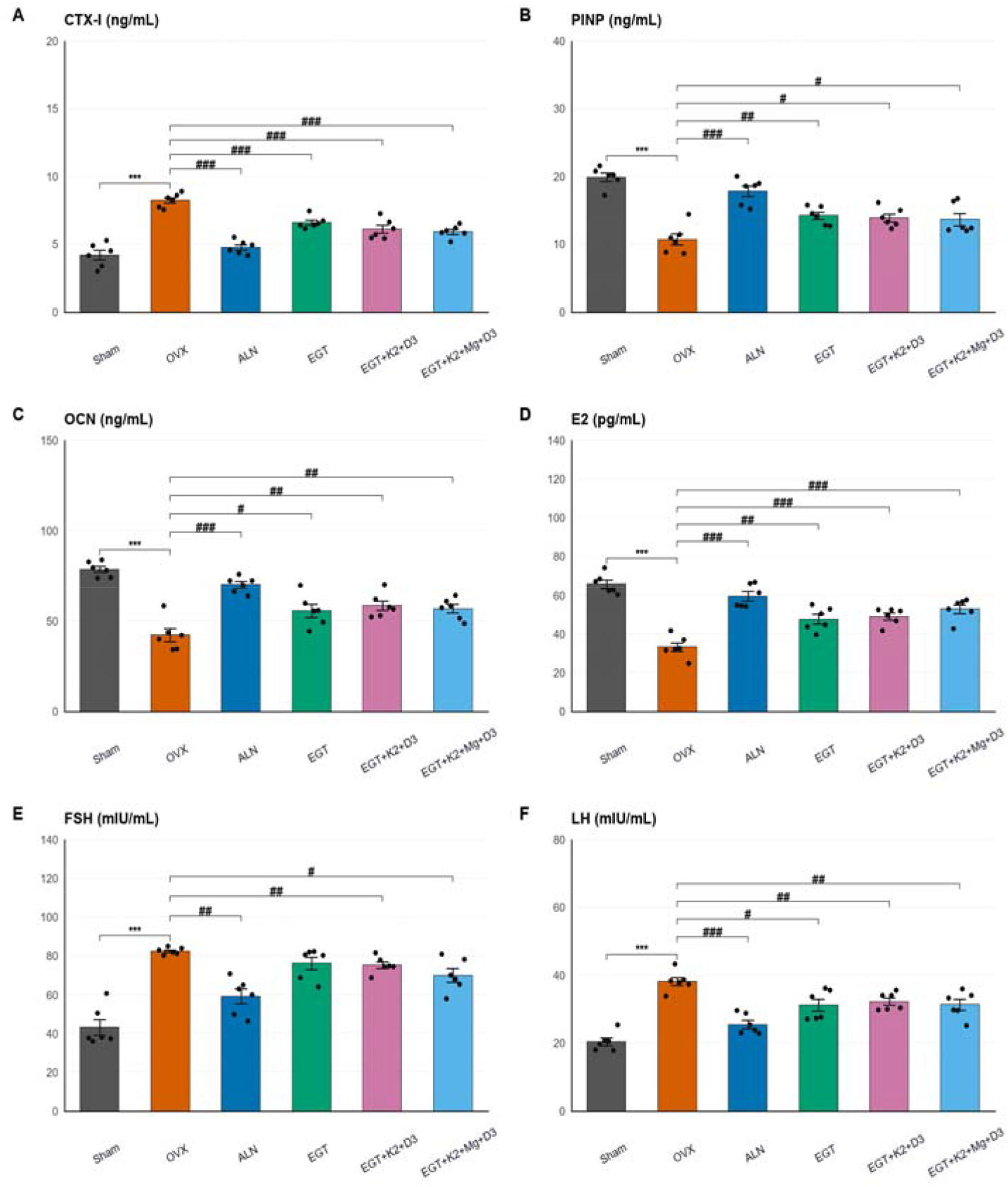
Serum bone turnover markers and sex hormone profile. OVX increased CTX-I and decreased PINP and osteocalcin/BGP. OVX also reduced E2 and increased LH and FSH. Alendronate and ergothioneine-based interventions reduced CTX-I, partially restored formation-associated markers, increased E2, and lowered LH.

Alendronate reduced CTX-I to 4.758 ± 0.500 ng/mL and raised PINP and osteocalcin/BGP to 17.84 ± 1.93 and 69.98 ± 4.43 ng/mL (all P < 0.01 vs OVX). EGT, EGT+K2/D3 and EGT+K2/MgT/D3 all significantly reduced CTX-I, to 6.585 ± 0.450, 6.094 ± 0.725 and 5.928 ± 0.449 ng/mL (all P < 0.01 vs OVX). PINP rose to 14.22 ± 1.32 (P < 0.05), 13.91 ± 1.43 (P < 0.05) and 13.64 ± 2.21 ng/mL (not significant), and osteocalcin/BGP to 55.66 ± 8.76 (P < 0.05), 58.49 ± 6.60 (P < 0.01) and 56.79 ± 5.75 ng/mL (P < 0.01), respectively. These changes indicate partial correction of the OVX-induced turnover imbalance, most consistently for CTX-I.

### Serum Sex Hormones (Exploratory)

OVX produced the expected endocrine phenotype: serum E2 decreased from 65.64 ± 5.09 (sham) to 33.20 ± 5.60 pg/mL (P < 0.01), whereas LH rose from 20.35 ± 2.73 to 38.17 ± 3.05 (P< 0.01) and FSH from 43.18 ± 10.02 to 82.46 ± 1.69 (P < 0.01). Alendronate increased E2 and lowered LH (both P < 0.01 vs OVX) and reduced FSH (P < 0.05 vs OVX) (Fig. 4).

EGT, EGT+K2/D3 and EGT+K2/MgT/D3 significantly increased E2, to 47.64 ± 5.92, 49.03 ± 4.31 and 52.77 ± 5.51 pg/mL (all P < 0.01 vs OVX), and lowered LH to 31.22 ± 4.35, 32.26 ± 2.55 and 31.32 ± 3.91 (all P < 0.05 vs OVX). FSH showed only non-significant numerical reductions (76.18 ± 7.93, 75.25 ± 4.24 and 69.99 ± 8.52; all P > 0.05). However, because bilateral ovariectomy removes the principal ovarian source of estradiol, the apparent rise in serum E2 cannot readily be ascribed to restored ovarian output and may partly reflect ELISA cross-reactivity or serum-matrix effects; these endocrine findings are therefore exploratory and require confirmation by LC-MS/MS-based steroid quantification.

### Serum Inflammatory and Oxidative Stress Markers

OVX markedly increased systemic inflammation: TNF-α rose from 414.0 ± 46.0 (sham) to 700.5 ± 27.4 pg/mL and IL-6 from 62.29 ± 5.89 to 121.2 ± 5.59 pg/mL (both P < 0.01). Alendronate lowered TNF-α and IL-6 to 503.0 ± 24.2 and 73.68 ± 9.64 pg/mL (both P < 0.01 vs OVX). EGT, EGT+K2/D3 and EGT+K2/MgT/D3 reduced TNF-α to 577.3 ± 67.2 (P < 0.05), 590.4 ± 84.0 (P < 0.05) and 556.7 ± 71.6 pg/mL (P < 0.01), and IL-6 to 94.48 ± 9.25, 91.80 ± 6.97 and 88.35 ± 11.97 pg/mL (all P < 0.01 vs OVX), confirming an anti-inflammatory effect (Fig. 5).

**Figure 5.**
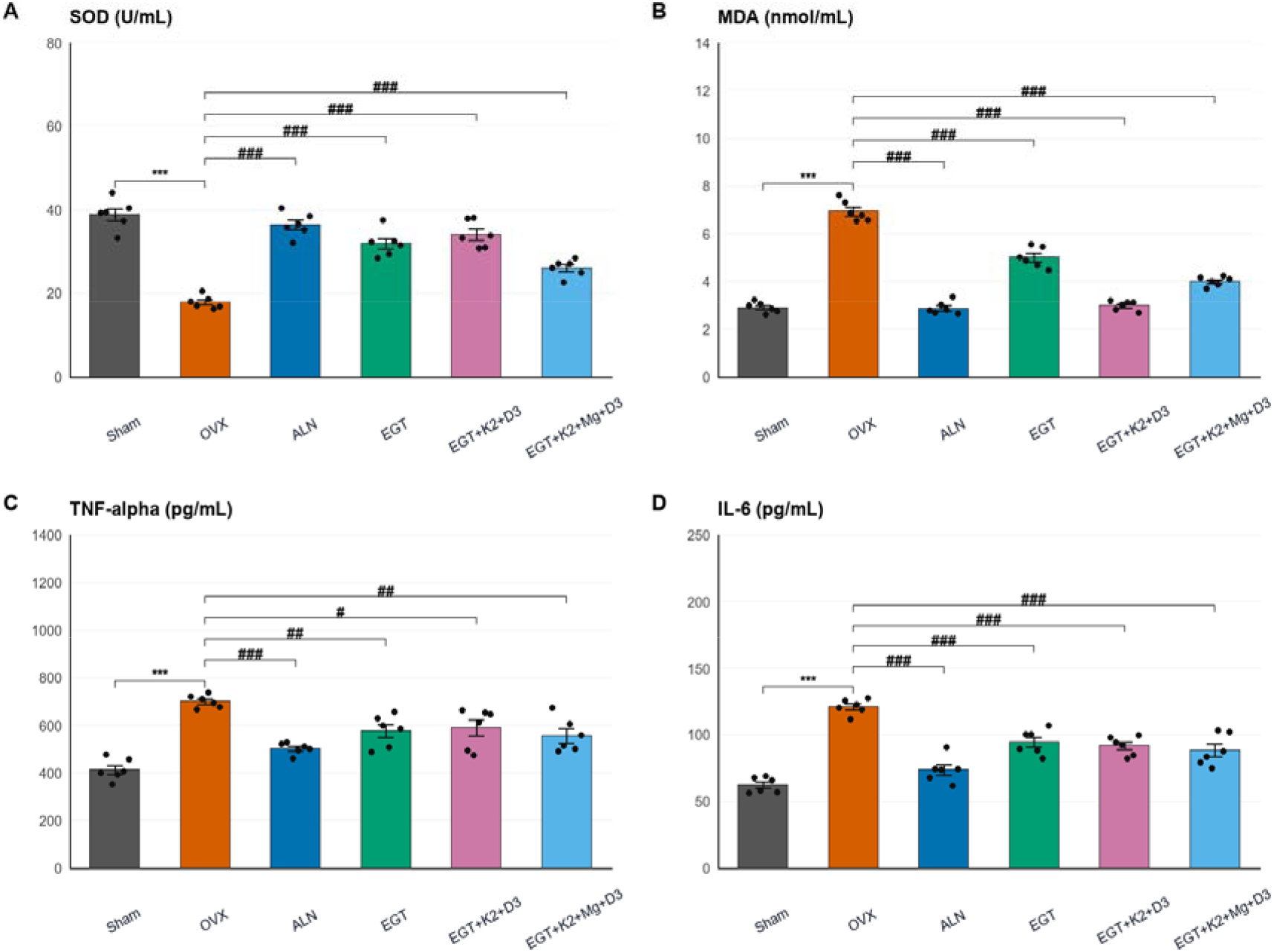
Inflammatory and oxidative stress markers. Ergothioneine-based interventions reduced TNF-α, IL-6, and MDA while increasing SOD activity relative to untreated OVX mice.

Oxidative-stress indices responded similarly. OVX decreased SOD from 38.91 ± 3.58 to 17.82 ± 1.51 U/mL and increased MDA from 2.903 ± 0.216 to 6.938 ± 0.431 nmol/mL (both P < 0.01). Alendronate restored SOD to 36.41 ± 2.82 U/mL and reduced MDA to 2.868 ± 0.266 nmol/mL (both P < 0.01 vs OVX). EGT, EGT+K2/D3 and EGT+K2/MgT/D3 increased SOD to 31.91 ± 3.16, 34.07 ± 3.27 and 26.05 ± 2.02 U/mL and reduced MDA to 5.009 ± 0.427, 2.985 ± 0.194 and 4.001 ± 0.210 nmol/mL (all P < 0.01 vs OVX). The strongest MDA suppression occurred in the EGT+K2/D3 group, whereas the magnesium-L-threonate-containing regimen showed a weaker SOD response than the other two (Fig. 5).

## Discussion

This study evaluated EGT, alone and combined with vitamin K2, vitamin D3, and magnesium L-threonate, in an OVX mouse model of bone loss. Across the systemic readouts examined–bone turnover, inflammatory cytokines, oxidative stress, and endocrine markers–the regimens produced consistent improvement, and distal-femoral trabecular area on histology rose significantly in every EGT-based group. Tibial micro-CT endpoints, by contrast, showed only non-significant trends, so the interventions did not meaningfully rebuild three-dimensional trabecular architecture under the conditions tested.

Model validity was clear. Uterine atrophy confirmed estrogen deficiency, and both micro-CT and histology demonstrated trabecular deterioration. The serum profile reinforced a high-risk skeletal state, with higher CTX-I, lower PINP and osteocalcin, elevated TNF-α and IL-6, reduced SOD, and increased MDA–a pattern that matches the established biology in which inflammatory and oxidative pathways activate osteoclasts and impair bone formation.

Alendronate behaved as a robust positive control, improving tibial BV/TV and Tb.N and normalizing CTX-I while partially restoring formation markers, confirming both model validity and assay sensitivity. The EGT-based regimens acted differently, with effects concentrated at the biomarker level: lower CTX-I in every group, variable increases in PINP and osteocalcin, and reduced TNF-α and IL-6. The accompanying fall in MDA and recovery of SOD point to antioxidant activity as a plausible contributor to the observed shifts in bone turnover.

An important caveat is that the histological and micro-CT endpoints were assessed at different skeletal sites—histology on the distal femur and micro-CT on the tibia—so they are not strictly comparable. The two modalities also sample different aspects of trabecular architecture and may differ in sensitivity depending on region selection, section plane, and thresholding. Treatment effects may have been sufficient to preserve local femoral trabecular area but insufficient to restore global tibial three-dimensional architecture. Finally, the analyzed sample size was modest and micro-CT variability was relatively large in some groups, so the study may have been underpowered to detect structural effects; larger, site-matched cohorts are needed to determine whether the favorable BV/TV, Tb.Th and Tb.N trends reach significance.

The endocrine results call for caution. Ergothioneine-based regimens raised E2 and lowered LH relative to OVX mice, with weaker effects on FSH. Because ovariectomy removes the main ovarian source of estradiol, the apparent rise in E2 is difficult to ascribe to restored ovarian output and may instead reflect assay variability, extra-ovarian steroid metabolism, altered clearance, or indirect endocrine effects. Confirmation will require orthogonal hormone assays, assessment of adrenal and adipose steroidogenesis, and measurement of estrogen-responsive tissue markers.

The combination design is an important constraint on interpretation. The study included an ergothioneine-alone arm and two progressively expanded combination arms, but no single-nutrient control arms (vitamin K2, vitamin D3, or magnesium L-threonate alone) and no full factorial structure. Consequently, although the data establish that the ergothioneine-based regimens are biologically active, they cannot attribute the effects to ergothioneine versus the co-administered nutrients, nor distinguish additive from synergistic interactions. Indeed, the responses of the EGT, EGT+K2/D3, and EGT+K2/MgT/D3 arms overlapped closely across most endpoints, providing no evidence that adding vitamin K2, vitamin D3, or magnesium L-threonate conferred benefit beyond ergothioneine alone; the apparent dose-dependent gradient in trabecular area was small and did not reach between-regimen significance. Resolving the individual and combined contributions will require single-component control arms and an adequately powered factorial design.

Several further limitations apply. Forty-eight animals were purchased to offset perioperative loss, and five to six per group were ultimately analyzed. Sham surgery was performed, but future studies should document that surgical stress was matched apart from ovarian removal. Micro-CT acquisition and region-of-interest parameters warrant fuller reporting in line with accepted guidelines, and the source, purity, and preparation of EGT, vitamin K2, vitamin D3, and magnesium L-threonate should be fully disclosed. Finally, mechanistic endpoints— biomechanical testing, dynamic histomorphometry, osteoclast and osteoblast staining, and pathway analyses—were not included and would strengthen causal claims.

In sum, EGT alone and combined with vitamin K2, vitamin D3, and magnesium L-threonate improved systemic bone-turnover, inflammatory, and oxidative-stress markers in OVX mice and preserved distal-femoral trabecular area, but did not significantly recover tibial micro-CT trabecular structure. The interventions therefore appear to act mainly on the pathological biological environment, offering limited structural protection under the conditions studied.

## Conclusions

EGT, alone and combined with vitamin K2, vitamin D3, and magnesium L-threonate, attenuated several systemic abnormalities of OVX, including increased bone resorption, impaired formation markers, inflammatory cytokine elevation, and oxidative stress; changes in sex-hormone measurements were also observed but remain exploratory pending assay confirmation. The regimens improved distal-femoral trabecular area, whereas recovery of tibial three-dimensional micro-CT structure was limited. These findings justify further study of EGT-based nutritional strategies for estrogen-deficiency-associated bone loss, incorporating mechanistic pathway analysis, biomechanical testing, dynamic histomorphometry, site-matched imaging, and dose optimization.

## Author Contributions

W.L. (Wei Liu), Y.T. (Yongxian Tang): conceptualization, methodology, formal analysis, writing—original draft. W.D. (Wei Ding): investigation, data curation. J.C. (Juan Cao): investigation, validation. C.G. (Cong Guo): formal analysis, visualization. G.X. (Guohua Xiao): conceptualization, supervision, project administration, writing—review and editing. All authors have read and agreed to the published version of the manuscript.

## Institutional Review Board Statement

The animal study protocol was approved under ethics approval number I20250006.

## Informed Consent Statement

Not applicable.

## Data Availability Statement

The data presented in this study are available from the corresponding author upon reasonable request. Processed analysis tables are available in the working dataset associated with this manuscript.

