## Supplementary_Materials for "Gene III® L-Ergothioneine, alone or combined with vitamin K2, vitamin D3 and magnesium L-threonate, attenuates bone turnover, inflammatory and oxidative disturbances in ovariectomized mice"

Supplementary tables for the manuscript: Ergothioneine Alone or Combined with Vitamin K2, Vitamin D3, and Magnesium L-Threonate Attenuates Bone Turnover Imbalance, Oxidative Stress, and Inflammation in Ovariectomized Mice.

**Supplementary Table S1. Experimental groups and treatments.**

| **Group** | **n analyzed** | **Animal IDs** | **Surgery** | **Treatment** | **Dose** |
| --- | --- | --- | --- | --- | --- |
| Sham control | 6 | Con-3 to Con-8 | Sham surgery | Water by oral gavage | Equivalent volume |
| OVX model | 6 | Model-3 to Model-8 | Bilateral ovariectomy | Water by oral gavage | Equivalent volume |
| Positive control | 6 | Gu-yang-1 to Gu-yang-6 | Bilateral ovariectomy | Alendronate | 1.53 mg/kg |
| Ergothioneine | 6 | A-3 to A-8 | Bilateral ovariectomy | Ergothioneine | 30 mg/kg/day |
| Ergothioneine + K2 + D3 | 6 | B-3 to B-8 | Bilateral ovariectomy | Ergothioneine + vitamin K2 + vitamin D3 | Ergothioneine 30 mg/kg/day + vitamin K2 40 µg/kg/day + vitamin D3 500 IU/kg/day |
| Ergothioneine + K2 + magnesium L-threonate + D3 | 6 | C-3 to C-8 | Bilateral ovariectomy | Ergothioneine + vitamin K2 + magnesium L-threonate + vitamin D3 | Ergothioneine 30 mg/kg/day + vitamin K2 40 µg/kg/day + magnesium L-threonate 350 mg/kg/day + vitamin D3 500 IU/kg/day |

**Supplementary Table S2. Micro-CT summary statistics.**

| **Endpoint** | **Unit** | **Sham control** | **OVX model** | **Alendronate** | **Ergothioneine** | **Ergothioneine + K2 + D3** | **Ergothioneine + K2 + magnesium L-threonate + D3** |
| --- | --- | --- | --- | --- | --- | --- | --- |
| BV/TV | % | 10.2 ± 2.17 | 5.58 ± 3.26 | 16.5 ± 5.82 | 5.92 ± 1.54 | 6.15 ± 2.36 | 6.37 ± 4.07 |
| Tb.N | 1/mm | 1.46 ± 0.378 | 0.708 ± 0.507 | 1.97 ± 0.785 | 0.976 ± 0.161 | 0.978 ± 0.260 | 0.986 ± 0.702 |
| Tb.Th | mm | 0.0744 ± 0.00347 | 0.0633 ± 0.00485 | 0.0769 ± 0.0107 | 0.0651 ± 0.00646 | 0.0676 ± 0.00683 | 0.0661 ± 0.00981 |
| Tb.Sp | mm | 0.252 ± 0.0394 | 0.327 ± 0.107 | 0.217 ± 0.0331 | 0.292 ± 0.0352 | 0.313 ± 0.0717 | 0.301 ± 0.0847 |
| Tb.Pf | 1/mm | 34.6 ± 2.62 | 41.4 ± 3.72 | 27.5 ± 4.34 | 39.5 ± 3.47 | 40.2 ± 6.01 | 39.7 ± 3.08 |
| SMI |  | 3.08 ± 0.195 | 3.39 ± 0.434 | 2.76 ± 0.101 | 3.16 ± 0.0948 | 3.25 ± 0.172 | 3.22 ± 0.185 |
| Conn.Dn | 1/mm^3 | 45.4 ± 15.6 | 26 ± 31.4 | 74.4 ± 33.3 | 15.7 ± 10.5 | 18.6 ± 9.95 | 18.8 ± 27.7 |

**Supplementary Table S3. Serum biochemical and ELISA summary statistics.**

| **Endpoint** | **Unit** | **Sham control** | **OVX model** | **Alendronate** | **Ergothioneine** | **Ergothioneine + K2 + D3** | **Ergothioneine + K2 + magnesium L-threonate + D3** |
| --- | --- | --- | --- | --- | --- | --- | --- |
| CTX-I | ng/mL | 4.19 ± 0.883 | 8.24 ± 0.512 | 4.76 ± 0.5 | 6.59 ± 0.45 | 6.09 ± 0.725 | 5.93 ± 0.449 |
| PINP | ng/mL | 19.9 ± 1.51 | 10.7 ± 2.11 | 17.8 ± 1.93 | 14.2 ± 1.32 | 13.9 ± 1.42 | 13.6 ± 2.21 |
| OCN | ng/mL | 78.6 ± 4.24 | 42.1 ± 8.87 | 70 ± 4.43 | 55.7 ± 8.76 | 58.5 ± 6.6 | 56.8 ± 5.75 |
| E2 | pg/mL | 65.6 ± 5.09 | 33.2 ± 5.6 | 59.5 ± 5.89 | 47.6 ± 5.92 | 49 ± 4.31 | 52.8 ± 5.51 |
| FSH | mIU/mL | 43.2 ± 10 | 82.5 ± 1.69 | 59.1 ± 9.32 | 76.2 ± 7.93 | 75.3 ± 4.24 | 70 ± 8.52 |
| LH | mIU/mL | 20.4 ± 2.73 | 38.2 ± 3.05 | 25.5 ± 2.95 | 31.2 ± 4.35 | 32.3 ± 2.55 | 31.3 ± 3.91 |
| SOD | U/mL | 38.9 ± 3.58 | 17.8 ± 1.51 | 36.4 ± 2.82 | 31.9 ± 3.16 | 34.1 ± 3.27 | 26.1 ± 2.02 |
| MDA | nmol/mL | 2.9 ± 0.216 | 6.94 ± 0.431 | 2.87 ± 0.266 | 5.01 ± 0.427 | 2.98 ± 0.194 | 4 ± 0.21 |
| TNF-alpha | pg/mL | 414 ± 46 | 701 ± 27.4 | 503 ± 24.2 | 577 ± 67.2 | 590 ± 83.9 | 557 ± 71.6 |
| IL-6 | pg/mL | 62.3 ± 5.89 | 121 ± 5.59 | 73.7 ± 9.64 | 94.5 ± 9.25 | 91.8 ± 6.97 | 88.4 ± 12 |

**Supplementary Table S4. Histological trabecular parameters.**

| **Group** | **n** | **Histological Tb.N (/mm)** | **Trabecular area (%)** | **Empty lacuna rate** |
| --- | --- | --- | --- | --- |
| Sham control | 6 | 3.833 ± 1.169 | 16.41 ± 4.496 | 0 |
| OVX model | 6 | 2.333 ± 0.5164 | 7.562 ± 2.615** | 0 |
| Alendronate | 6 | 3.833 ± 1.169 | 16.90 ± 1.516## | 0 |
| Ergothioneine | 6 | 3.000 ± 1.095 | 14.19 ± 4.832# | 0 |
| Ergothioneine + K2 + D3 | 6 | 2.833 ± 0.7528 | 14.55 ± 3.554# | 0 |
| Ergothioneine + K2 + magnesium L-threonate + D3 | 6 | 2.667 ± 0.8165 | 14.95 ± 4.462# | 0 |
